# Discovery of an endogenous DNA virus in the amphibian killing fungus and its association with pathogen genotype and virulence

**DOI:** 10.1101/2023.03.16.532857

**Authors:** Rebecca Clemons, Mark Yacoub, Evelyn Faust, L. Felipe Toledo, Thomas S. Jenkinson, Tamilie Carvalho, D. Rabern Simmons, Erik Kalinka, Lillian K. Fritz-Laylin, Timothy Y. James, Jason E. Stajich

**Affiliations:** Department of Ecology and Evolutionary Biology, University of Michigan, Ann Arbor, MI 48109, USA; Department of Microbiology and Plant Pathology, University of California Riverside, Riverside, CA 92521, USA; Laboratório de História Natural de Anfíbios Brasileiros (LaHNAB), Departamento de Biologia Animal Instituto de Biologia (IB), Universidade Estadual de Campinas, Campinas, SP, 13083-862, Brazil; Department of Biological Sciences, California State University, East Bay, Hayward, CA 94592, USA; Department of Biology, University of Massachusetts Amherst, Amherst, MA 01003 USA

**Keywords:** Mycovirus, Chytrid, *Batrachochytrium*, CRESS Virus, *Circoviridae*, Parasite, Amphibian disease

## Abstract

The Global Panzootic Lineage (GPL) of the pathogenic fungus *Batrachochytrium dendrobatidis* (*Bd*) has caused severe amphibian population declines, yet the drivers underlying the high frequency of GPL in regions of amphibian decline are unclear. Using publicly available *Bd* genome sequences, we identified multiple non-GPL *Bd* isolates that contain a circular Rep-encoding single stranded DNA (CRESS)-like virus which we named BdDV-1. We further sequenced and constructed genome assemblies with long read sequences to find that the virus is integrated into the nuclear genome in some strains. Attempts to cure virus positive isolates were unsuccessful, however, phenotypic differences between naturally virus positive and virus negative *Bd* isolates suggested that BdDV-1 decreases the growth of its host *in vitro* but increases the virulence of its host *in vivo*. BdDV-1 is the first described CRESS DNA mycovirus of zoosporic true fungi with a distribution inversely associated with the emergence of the panzootic lineage.

## Introduction

Recent observations of the decline in global amphibian diversity coincide with population genetic evidence of the global spread and emergence of the fungal pathogen *Batrachochytrium dendrobatidis* (*Bd*) ^1–4^. The pathogen causes the disease chytridiomycosis in susceptible amphibian species through heavy skin infection that can cause mortality due to osmolyte imbalance and electrolyte depletion^5^. *Bd* is a generalist pathogen and is associated with the decline of over 500 amphibian species, with most of these declines attributed to the invasion of a low diversity, rapidly expanded Global Panzootic Lineage of *Bd* (*Bd*-GPL)^4,6,7^. Other *Bd* lineages (such as *Bd*-BRAZIL and *Bd*-CAPE) are mostly enzootic to limited geographic regions and generally do not cause large amphibian declines as has been observed with *Bd*-GPL^6,8,9^. Yet, the drivers that have facilitated the recent expansion of *Bd* and *Bd*-GPL are currently unknown. One possibility is that *Bd* has escaped from its natural enemies, such as hyperparasitic viruses or other fungi. Release from population control by natural enemies is hypothesized to be behind the recent emergence of other emergent fungal diseases^10,11^. However, no such enemies of *Bd* have been found to date. Here we describe the discovery of a novel DNA mycovirus of *Bd* recovered almost exclusively from the less virulent non-GPL *Bd* lineages.

Mycoviruses, which most commonly have a dsRNA genome, are common hyperparasites of fungi and are known to impact host fitness and virulence^12,13^. Perhaps the best known example of a mycovirus that impacts host virulence is CHV1, which infects the agent of chestnut blight, *Cryphonectria parasitica*. CHV1 reduces the virulence of its host and has been used with limited success as a biocontrol agent against chestnut blight in some parts of Europe^14^. *Bd* is a member of the Chytridiomycota, a phylum that is primarily aquatic and reproduces with motile zoospores. There is minimal knowledge about the prevalence and diversity of mycoviruses in the zoosporic lineages of fungi. Previous attempts to screen *Bd* isolates for the presence of dsRNA mycoviruses have yielded negative results^15,16^. Recently, mycoviruses with small ssDNA genomes related to Circular Rep-Encoding Single Stranded (CRESS) DNA viruses have been described from Ascomycete fungi and shown to have a negative impact on host virulence^17–19^. The characterized CRESS viruses in fungi are members of family *Genomoviridae*, however, all characterized viruses infect Ascomycota, yet there is genomic evidence from endogenized CRESS virus genes that the distribution of this group is phylogenetically more widespread in fungi^20^. This demonstration that CRESS viruses are more common than appreciated in fungi^20^ prompted us to screen for similar mycoviruses in *Bd.* Upon the discovery of such a virus in multiple isolates, we tested for the potential impact of these viruses on fungal phenotypes including virulence towards a model amphibian host.

## Results

### Viral Discovery

To test how common CRESS viruses are in *Bd*, we screened publicly available *Bd* strain genome sequences from all existing lineages^4,6,21^ for DNA mycoviruses by *de novo* assembling reads that did not align to the reference genome JEL423. We found that numerous isolates contain viral genes — Rep, encoding a replication-associated protein, and Cap, encoding a capsid protein — indicative of a CRESS DNA mycovirus infection. These virus positive isolates almost exclusively lie within less virulent enzootic *Bd* lineages (**Figure 1A**): Despite *Bd*-GPL being the most sampled lineage, only 1 out of 211 sequences had matches to viral genes (**Figure 1A**). The enzootic *Bd*-BRAZIL lineage is distinctive because less than half of the genome data from these isolates are positive for CRESS virus genes. The other enzootic lineages of *Bd-*ASIA1 and *Bd*-CAPE have a higher proportion of isolates with copies of the viral genome, in some cases up to 100% of screened isolates. Notably, the copy number of the virus relative to the genome average varied between strains (ranging from 0.08-2.49) and was lower in *Bd*-BRAZIL and *Bd*-CAPE compared to *Bd*-ASIA1 (**Figure 1B**). Additional DNA and protein sequence searches of available *B. salamandrivorans*, the sister species of *Bd*, genomic data did not identify any CRESS-like viral sequences.

**Figure 1:**
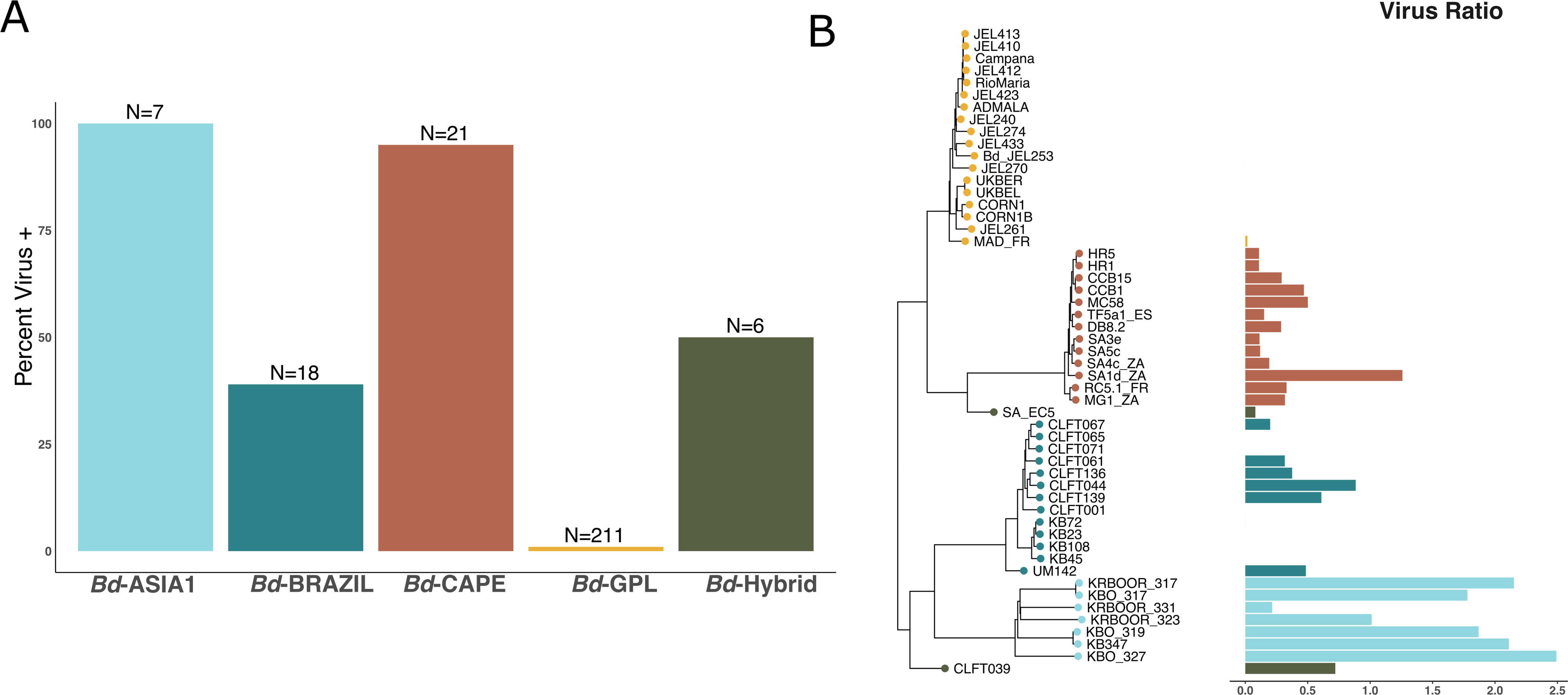
BdDV-1 is mostly absent in *Bd*-GPL. **A)** Barplot showing percent of infected strains of sampled *Bd* lineages. N indicates the number of genome sequence isolates analyzed. **B)** Phylogeny of *Bd* strains constructed from Single Nucleotide Polymorphisms (left; x-axis indicates genetic distance based on SNPs) and barplot showing the ratio of Virus to Fungus sequence reads found in each genome sequencing set (right), colored by *Bd* lineage.

### Viral Phylogeny and Assembly

We have named this mycovirus Batrachochytrium dendrobatidis DNA Virus 1 (BdDV-1). The viral phylogeny constructed from the Rep protein places BdDV-1 in the family *Circoviridae* (**Figure 2A**). The BdDV-1 lineage is distinct from other identified fungal CRESS viruses and has a sister relationship to those recovered from environmental sequencing of water, sewage, or viruses associated with animals. The structure of the BdDV-1 genomes recovered from *Bd* isolates has three or four Open Reading Frames (ORFs) depending on the strain, including *Cap* and *Rep* encoding ORFs that are ubiquitous in other CRESS viruses. Comparing the phylogeny of BdDV-1 with a dendrogram of *Bd* strains shows a general fidelity of the virus between *Bd* lineages (Baker’s Congruency = 0.94), although there appears to be little co-phylogeny within individual lineages (**Figure 2B**). Interestingly, the *Bd-*GPL strain MAD-FR appears to have acquired its BdDV-1 infection from the *Bd*-CAPE lineage that has been introduced into Europe^4,21^.

**Figure 2:**
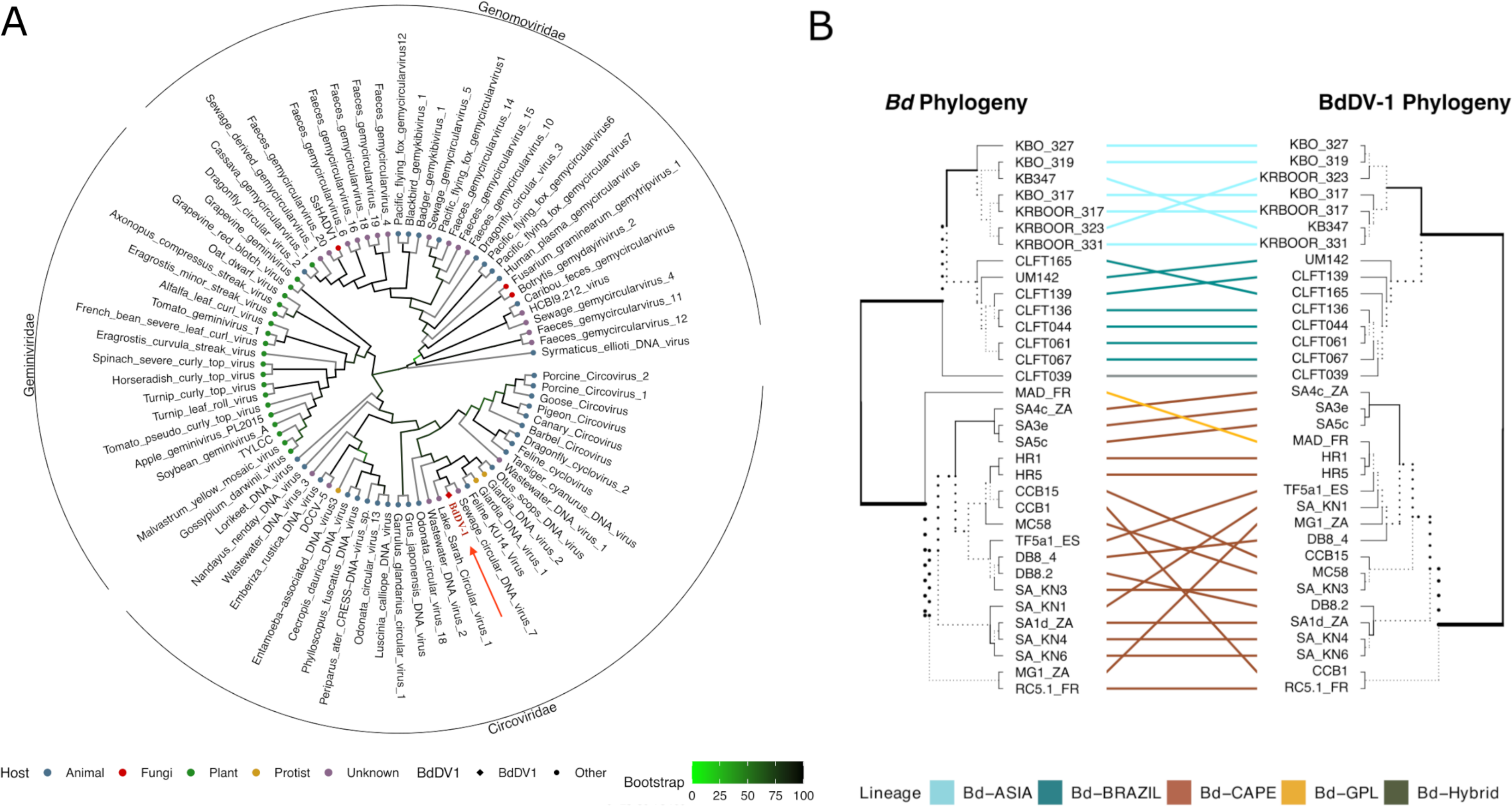
BdDV-1 belongs to the family *Circoviridae* and shares a close evolutionary relationship with *Bd*. **A)** A cladogram of the Maximum Likelihood tree estimated from Rep amino acid sequence of BdDV-1 show it as a member of family *Circoviridae* and distinct from and other mycoviruses (red). Tips are color coded based on viral hosts and branches are colored by SH-like support values. BdDV-1 is highlighted with a red arrow. **B)** A tanglegram demonstrating co-phylogeny between infected *Bd* strains (left) and BdDV-1 (right). BdDV-1 ML phylogenetic tree constructed based on nucleotide sequences of the *Cap* gene. Branch weight indicates common sub-branches between the two trees. See also Figure S2.

The Illumina-based genome assemblies of the virus produced 2-4 kb linear viral contigs with some direct repeats and overlap in the assembly graph indicative of a possible circular genome (**Figures S1A-D**). Assuming the repeats could represent a circular viral genome, then the genome would be 2.2 kb in length with a single capsid gene and a replication-associated protein gene **(Figure S1C**), however lacking the canonical stem-loop motif associated with a nonanucleotide that is characteristic of CRESS viruses and required for replication^22^. PCR with outward-facing primers was consistent with the presence of a circular form of the virus (**Figure S1B**). However, attempts to amplify the circular version of the virus using a modified rolling circle amplification protocol^23^ using multiple approaches were unsuccessful. These data suggest that if there is a circular form it should be rare, not the dominant structure of the BdDV-1 genome, and unable to undergo normal rolling circle replication due to lack of appropriate motifs.

### Viral Integration

To test whether BdDV-1 infection impacts the phenotype of its host, as is the case with other DNA mycoviruses^17–19^, we attempted to generate a virus-free isolate using antivirals. A paired infected and virus-cured *Bd* isolate would allow for the direct comparison of the effects of BdDV-1 on host fungal phenotypes. We tested 71 single zoosporangia isolations from a virus positive *Bd* isolate (CLFT139) grown in control media, or in presence of the antivirals cycloheximide or ribavirin. As each zoosporangium develops from a single zoospore, a single zoosporangium isolate should derive from a single cell. However, all these curing attempts were unsuccessful, as determined by a PCR test for the presence of the *Rep* gene after subculturing.

Due to the inability to clear the virus, the failure to amplify the genome via rolling circle replication, and low coverage of the virus in the genome sequencing data (**Figure 1B**), we investigated whether the virus may integrate into the genome. We achieved this by sequencing multiple virus-infected strains using long read Oxford Nanopore Technology (ONT). Our assembled *Bd*-BRAZIL isolate CLFT044 genome contained a 4.405 kb locus with the BdDV-1 genome integrated adjacent to a rDNA locus in a sub-telomeric region of scaffold_10. The total viral locus is 4.4kb long and encodes four putative ORFs, a *Cap* and *Rep* gene followed by ORF3 and ORF4 which are homologs of a *Rep* gene. ORF3 and ORF4 are similar to Rep proteins from other CRESS viruses, but are both phylogenetically distinct from BdDV-1 Rep protein (**Figure S2**). Additionally, ORF3 and ORF4 shared nearly 100% sequence similarity with each other indicating they may have originated from a duplication or partial integration event.

The viral genome was integrated into a locus 21.2 kb proximal to the telomere, upstream of a GAG-pre integrase gene, and downstream of a rDNA sequence (**Figure 3A**). Although the ∼1 Mb scaffold where BdDV-1 integrated is shared among *Bd* strains, the 21.12 kb region between the telomere and BdDV-1 locus was not found in virus negative (v-) strains, including the reference genome, JEL423. This subtelomeric region in CLFT044 did not contain any predicted protein coding genes (**Figure 3B)**. We used PCR primers targeting the rDNA region to the BdDV-1 *Rep* ORF in viral positive *Bd*-BRAZIL strains (CLFT044, CLFT061, and CLFT067) and viral negative strains (CLFT071, CLFT085, and JEL423). All viral positive strains also tested positive for the same integration locus (adjacent to the rDNA sequence).

**Figure 3:**
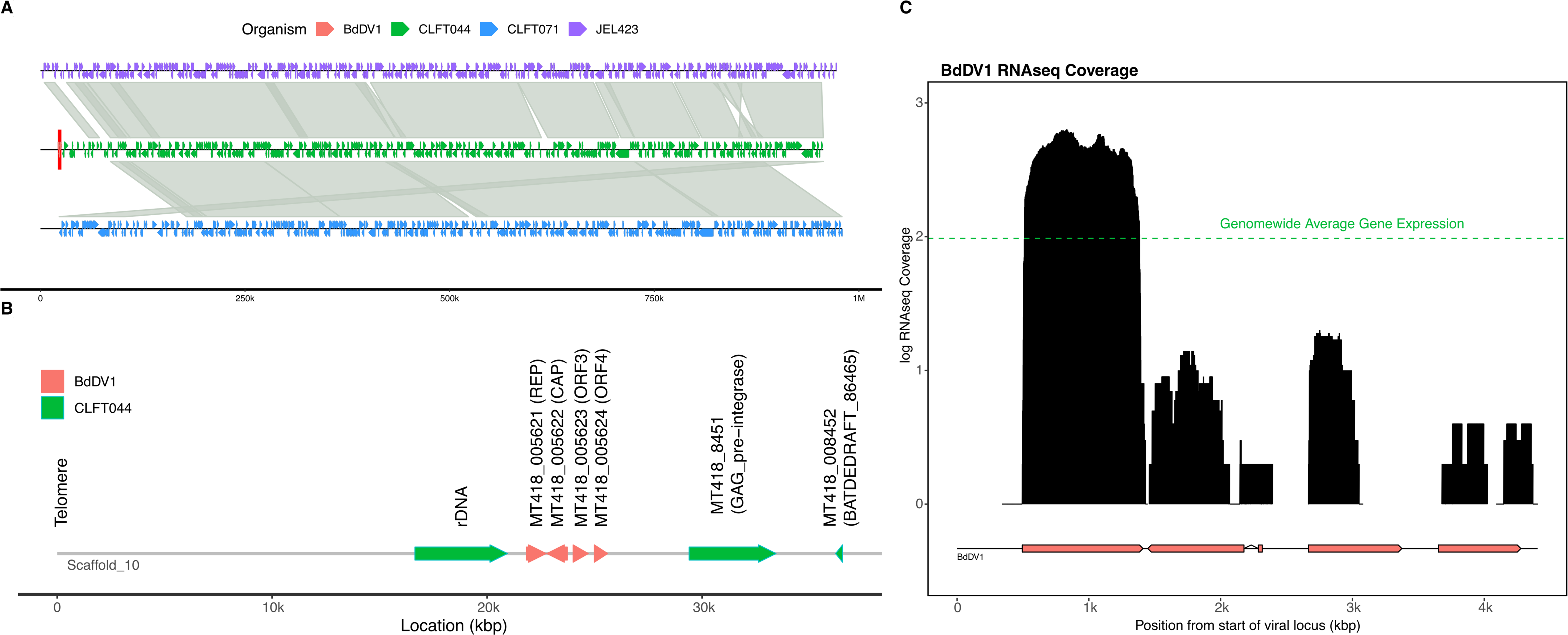
BdDV-1 is integrated into the *Bd* genome and remains transcriptionally active. **A)** View of integration supercontigs from 3 *Bd* strains: v+ strain, CLFT044 (green) and v-strains, CLFT071 (blue) and JEL423 (violet). Shaded linkage shows regions of synteny between the scaffolds. The BdDV-1 integration site is indicated in the CLFT044 scaffold by red shading. **B)** BdDV-1 integration locus in CLFT044 at the end of scaffold 10, including immediate downstream genes (green arrows). Locus IDs in CLFT044 annotation and Functional annotations are indicated. Genes of unknown function are labeled with their highest BLAST hit in reference genomes JEL423 or JAM81. The region spanning the telomere to ∼100kb region is not shared between infected and uninfected strains. **C)** BdDV-1 locus structure with log transformed expression at each position indicated using black bars. The average log transformed expression across all *Bd* genes is depicted by the green horizontal line. The *Rep* gene alone is highly expressed compared to fungal genes.

To further explore the localization of BdDV-1, we designed FISH probes to detect the viral genome, as well as a probe to the actin gene as an endogenous *Bd* gene as a control. We used these probes to localize BdDV-1 and actin genes in both v-JEL423 and v+ CLFT044 (**Figure 4A**). We quantified the number of hybridization signal spots and found that the actin control probe localized to either one or two spots in 58% of JEL423 and 32% of CLFT044 cells, while the BdDV-1 probe localized to only 3% of JEL423 cells but 46% of CLFT044 cells (**Figure 4B**). The vast majority of these localization spots co-localized with nuclear DNA signal. The number of foci and the localization of the BdDV-1 compared to the actin gene are consistent with chromosomal integration of the BdDV-1 virus in CLFT044. A BdDV-1 localization primarily within the *Bd* genome would explain why attempts to cure the virus have been unsuccessful. We also compared the subcellular structure of zoospores from a v+ and a v-strain using transmission electron microscopy. We did not detect obvious virions in the cytoplasm or nucleus of the v+ strains, nor did we observe changes to the organization of the cell (**Figure S3**), again, consistent with an endogenous localization of BdDV-1.

**Figure 4:**
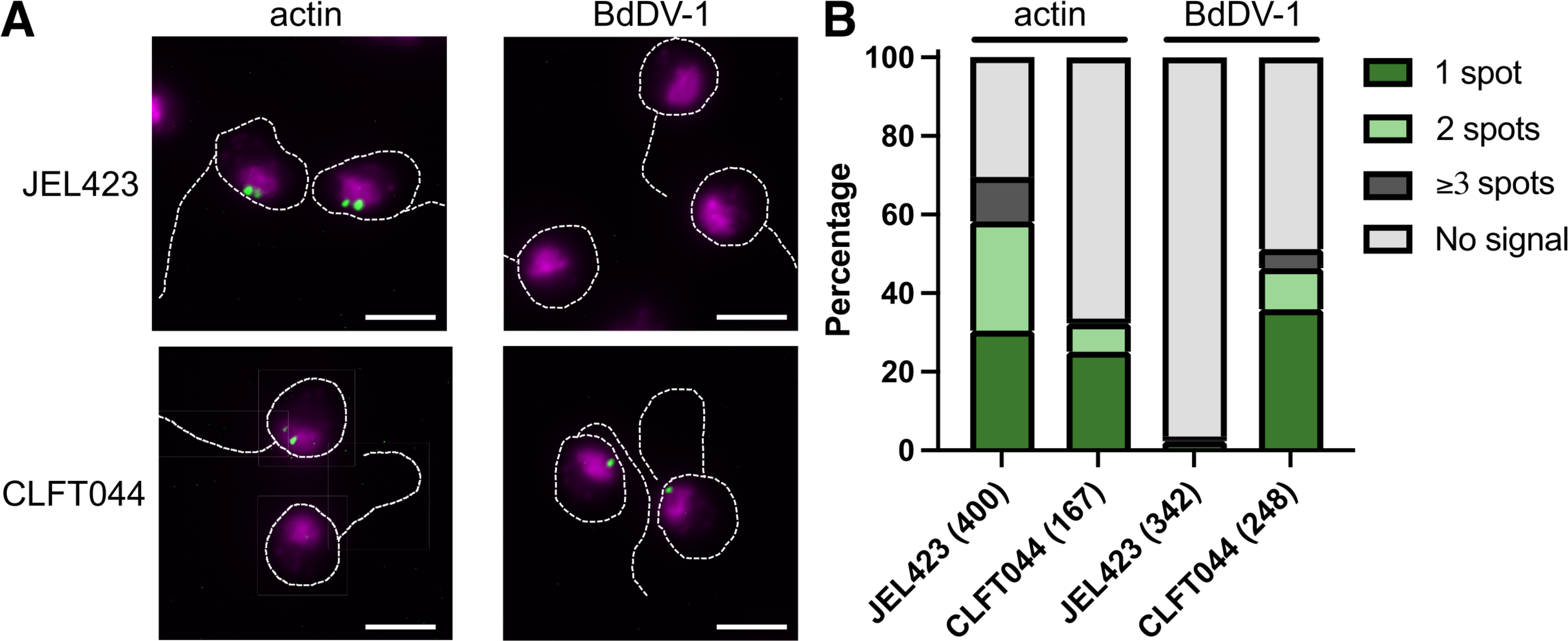
BdDV-1 genome localization using DNA FISH. **A)** Zoospores from v-strain JEL423 and v+ strain CLFT044 were fixed and stained with DNA probes targeting the *Bd* actin locus (left) or a region of the BdDV-1 genome (right). Images show overlays of DNA probe (green) and DAPI (magenta) fluorescence, dashed lines indicate cell boundaries and flagella. The brightness and contrast of probe signal were increased in the actin/CLFT044 image to improve visibility of foci. Scale bars: 5 µm. **B)** Quantification of hybridization signal spots per cell. Numbers of cells analyzed are indicated in parentheses.

To determine whether BdDV-1 ORFs are expressed, we extracted and sequenced mRNA from the v+ strain CLFT044 and found RNASeq reads that matched the viral ORFs (**Figure 3C**). The *Rep* gene was the most transcriptionally active viral gene and more highly expressed than the average expression observed in all *Bd* genes.

To identify differentially expressed genes (DEGs) between v+ and v-*Bd*-BRAZIL, we extracted and sequenced mRNA from three samples of the v+ strain, CLFT067, and three samples of the v-strain, CLFT071, all grown in 1% tryptone media. These strains were selected due to their close phylogenetic relationship and their shared geographic location^4,6,7^. We identified 97 DEGs between CLFT067 and CLFT071; among them, 46 DEGs were up-regulated in the v+ strain. Additionally, we detected multiple M36 domain-containing genes, putative virulence genes of *Bd*, exhibiting decreased expression in the v+ strain (**Figure S4**).

### Growth Assays of Infected Fungal Host

Because we were unable to generate isogenic v+ and v-isolates, we took advantage of the variation within the *Bd*-BRAZIL lineage to determine how the virus may impact its host. We hypothesized that since the *Rep* gene was present in primarily less virulent non-GPL lineages, BdDV-1 could reduce the virulence of *Bd* strains. The *Bd*-BRAZIL lineage contains a mixture of isolates, some carrying BdDV-1, as detected in genome sequences and validated by PCR of the *Rep* gene. We selected 8 phylogenetically distinct isolates of *Bd*-BRAZIL — 4 naturally v+ and 4 naturally v- (**Figure 1B**) — for growth and experimental infection assays.

We conducted two experiments to measure *in vitro* growth differences between v+ and v- *Bd* isolates. The first experiment consisted of three growth curve optical density assays that differed only in the total number of isolates and replicates. In all assays, zoospores were harvested from v+ and v- *Bd* isolates and diluted at equal concentrations into fresh media. Optical density (OD) readings were recorded for each isolate and averaged between replicates for the duration of the growth period (**Figure 5A**). In all three assays, the growth rate was not different between v+ (1: *M*=1.73, *SD*=0.44, 2: *M*=0.47, *SD*=0.25, 3: *M*=1.40, *SD*=0.11) and v- (1: *M*=2.37, *SD*=1.34, 2: *M*=0.71, *SD*=0.34, 3: *M*=1.25, *SD*=0.14) isolates, *t*(6)=0.87, *p*=0.416, *t*(4)=0.95, *p*=0.396, *t*(4)=1.71, *p*=0.163 for assays 1, 2 and 3 respectively (**Figure 5B**). However, the efficiency of growth (EOG) or carrying capacity, measured as the difference between the endpoint OD and initial OD, was higher among v- (1: *M*=0.19, *SD*=0.06, *M*=0.12, *SD*=0.04) compared to v+ (1: *M*=0.11, *SD*=0.02, 2: *M*=0.08, *SD*=0.02) isolates for the first two assays, *t*(6)=4.22, *p*<0.001 and *t*(4)=3.33, *p*=0.029 for assays 1 and 2, respectively (**Figure 5B**). There was no significant difference in EOG between v+ (*M*=0.11, *SD*=0.03) and v- (*M*=0.11, *SD*=0.03) isolates in the third assay, t(4)=0.42, p=0.698. In our second experiment, we inoculated media with zoospores of v+ and v- strains and tracked four replicates over 16 days using qPCR of the ITS region to measure *Bd* cell numbers, or zoospore equivalents (ZSE). Due to natural variation in ITS copy number among the strains used in this study, which ranged from 28-56 copies per genome, we adjusted growth curves to account for differences in relative ribosomal RNA copy number. Growth peaked at day 7 of day 10 for most strains (**Figure 5C**). Differences in ZSE were significant at day 7, with v- strains reaching a higher peak than v+ strains, (*M_v-_*=8.5e+05, *SD_v-_*=2.6e+05, *M_v+_*=3.8e+05, *SD_v+_*=2.4e+05, *t*(6)=2.65, *p*<0.05; **Figure 5D**). The relative ratio of fungal ITS rDNA (ZSE) to viral *Rep* gene copies was also monitored in experiment 2 for the v+ strains. We found that ratios of Virus/ZSE varied between 0.4-11.2, with a median of 1.1 (**Figure S5**). These results indicate that BdDV-1 is stably present and impacts the growth of *Bd in vitro* by reducing cell density at carrying capacity in both optical density and qPCR experiments.

**Figure 5:**
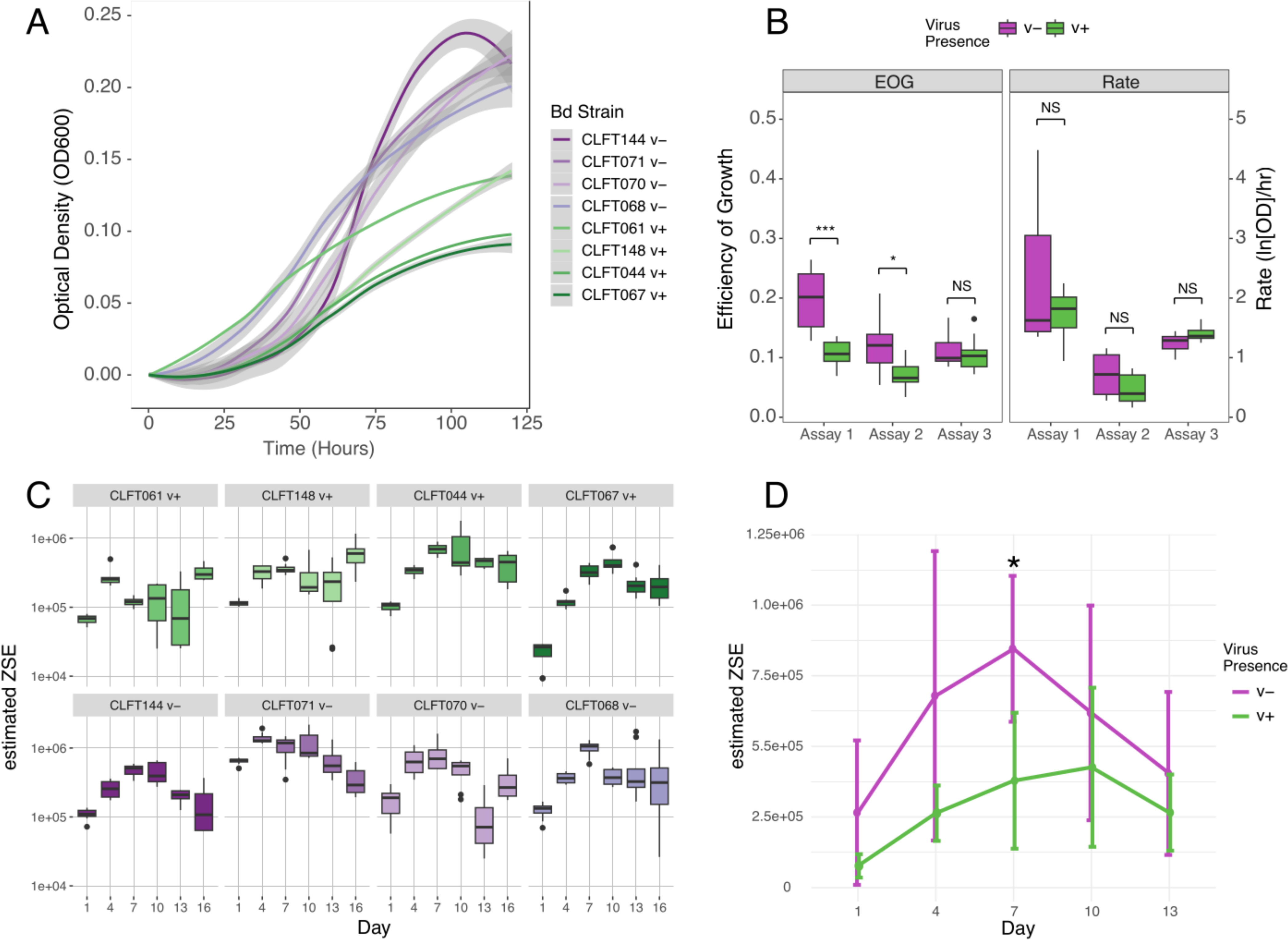
*Bd* isolates naturally infected with BdDV-1 (v+) show reduced growth efficiency compared to virus-free strains (v-). **A)** Growth curves are shown for infected (v+, green) and uninfected (v-, purple) strains. Gray shading indicates the 95% confidence interval. **B**) Comparisons of averages between all v- and v+ isolates for efficiency of growth (EOG) and rate of growth over three independent assays. **C**) Growth of infected (v+, green) and uninfected (v-, purple) strains in vitro estimated using qPCR. Values are shown in zoospore equivalents (ZSE), adjusted for differences in ribosomal ITS copy numbers using genome sequences. **D**) Comparison of growth curves at 5 time points estimated by qPCR aggregated across v+ (green) and v- (purple) strains. Bars indicate 1 standard deviation. Significant differences between groups are indicated as * = (*p*<0.05) and *** = (*p*<0.001). See also Figure S5.

### Experimental Infection of Amphibians

To determine the effects of BdDV-1 on *Bd* virulence we infected Dwarf Clawed frogs (*Hymenochirus boettgeri*) and monitored survivorship, *Bd* load in ZSE, and BdDV-1 copies per zoospore. Nine groups of 10 frogs each were individually infected with zoospore suspensions from 4 v+ Bd-BRAZIL isolates, 4 v- *Bd*-BRAZIL isolates, or received no infection as a control. The frogs were monitored daily for mortality and morbidity for the duration of the 60 day experiment. All 40 frogs infected with v+ isolates died of their infections by day 43. In contrast, only 28 of 40 frogs infected with v- isolates died of their infection by the end of the 60 days. All 10 control frogs and 12 of 40 frogs infected with v- isolates survived the experiment (**Figure 6A**). Only one v- *Bd* isolate (CLFT144) had 100% mortality. Despite this outlier, we tested for an effect of BdDV-1 on virulence using a Cox proportional hazards model. Frogs infected with v+ isolates had a significantly higher risk of death compared to those infected with v- isolates (HR=6.81, 95% CI 1.74-26.64, *p*=0.006). There was no correlation between initial body size and survival (*r*(78)=0.022, *p*=0.847). These results indicate that, contrary to our expectations, BdDV-1 is associated with increased *Bd* virulence.

**Figure 6:**
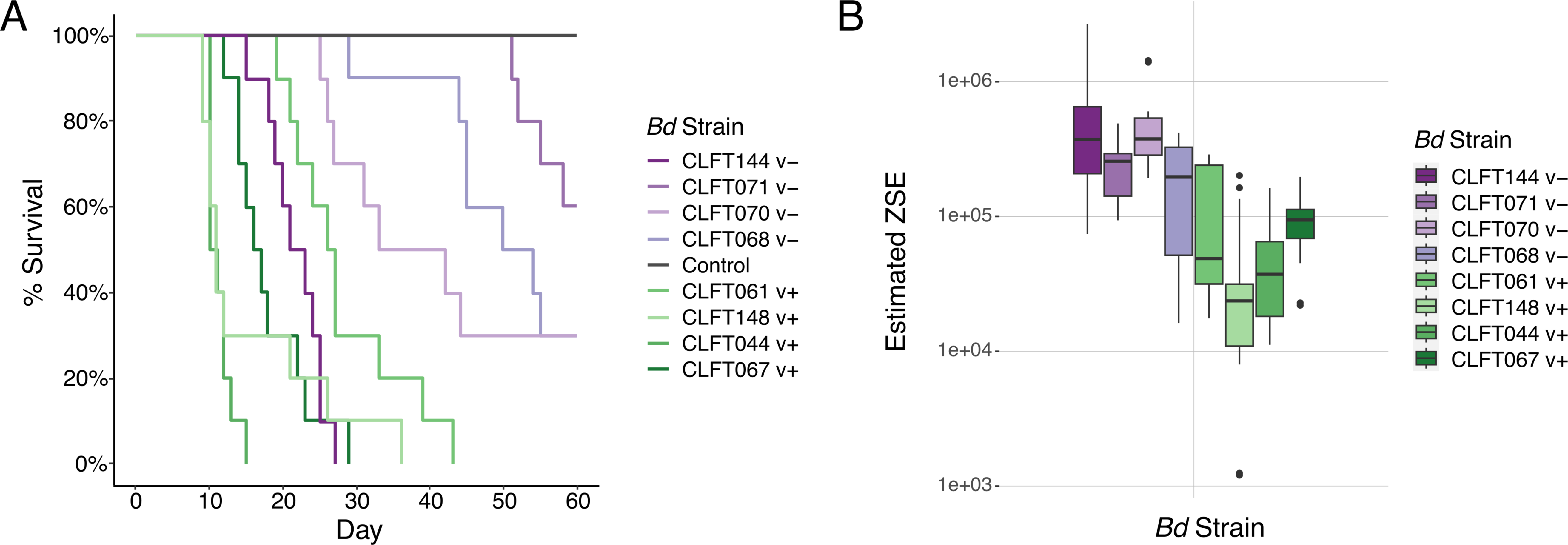
Frogs infected with v+ isolates have higher mortality rates but lower zoospore loads at death. **A)** Frog host survival for the 60 day experimental infection. Frogs infected with virus positive *Bd* strains (green) have a higher chance of mortality compared to frogs infected with virus negative *Bd* strains (purple). There was no mortality in the control group (gray). **B)** qPCR data showing the zoospore quantity (in zoospore equivalents ZSE) at time of death for all frogs that died during the experiment. Frogs infected with virus negative strains (purple) had higher zoospore loads at death. ZSE values were adjusted to account for different numbers of ribosomal ITS copy numbers as determined from genome sequences. See also Figure S6.

During the experiment, living frogs were also skin swabbed to monitor *Bd* zoospore load via qPCR on day 10, 30, and 60 post *Bd* exposure. A final skin swab was also taken at the time of death. For living frogs, high early time point zoospore loads correlated to higher probabilities of mortality during the experiment (**Figure S6A**). Among frogs that succumbed, those infected with v- isolates had a higher zoospore load at death (*M*=4.16×10^5^, *SD*=5.16×10^5^) compared to those infected with v+ isolates (*M*=7.66×10^4^, *SD*=7.53×10^4^), whether adjusting or not for differences in ITS copy number among strains, *t*(6)=3.54, *p*=0.01 (**Figure 6B**) and *t*(6)=5.5, *p*<0.001) (**Figure S6B**), respectively. The number of zoospores at death was not related to survival time, as individuals infected with the most virulent v- isolate (CLFT144) had similar mean zoospore loads at death to the least virulent v- isolate (CLFT071). This suggests that v+ *Bd* isolates may be associated with a higher virulence than v- isolates for an equivalent infection burden.

To determine the relationship between viral titer and infection outcome, we monitored the relative number of BdDV-1 copies per infected animal via qPCR on collected skin swabs. We found that viral titer, as measured by the ratio of viral copies to fungal nuclear genome copies, varied over time during the infection. After 10 days of infection, the ratio of viral copies to fungal nuclear genome copies was one or higher, and could be much higher (**Figure S6C**). For example, for strain CLFT061, the mean ratio of virus:fungus nuclear copies was 65.1. For all 4 v+ *Bd* isolates, the ratio decreased among living frogs from day 10 to day 30 post exposure and was lowest among deceased frogs (**Figure S6D**). This may indicate that while viral presence is significant for the relative virulence of the isolate, it is not necessarily the accumulation of viral genomes in these isolates that leads to increased virulence. Additionally, while virus to fungus ratios did vary between v+ isolates, it is unclear how this relates to virulence. It is notable that CLFT044 had the highest mortality rate and the lowest ratio of virus to fungus (**Figure S6C**). Causes and consequences of BdDV-1 variation in apparent titer are unclear and an area needing further work.

## Discussion

BdDV-1 is the first full length (containing both *Rep* and *Cap* genes) CRESS virus in the *Circoviridae* known to infect fungi. The cophylogeny and genome sequencing data suggest the viral sequence has a deep history mirroring the diversification of the fungus which may be explained by an endogenization event into the *Bd* genome. Although genome sequence data had been present for the strains used in these studies for over a decade, the presence of BdDV-1 had been overlooked for two reasons. First, most efforts to study mycoviruses have focused on RNA viruses, but DNA mycoviruses may be more prominent than realized^15,20^. Second, analysis which only maps DNA sequence to a single reference genome limits examination to elements found in the assembly; a pangenome approach for the diverse *Bd* lineages is needed to better describe the total diversity of genetic elements of the chytrid fungus^24,25^.

Our results suggest that BdDV-1 may be associated with decreased growth of its host *in vitro* but may also be associated with increased virulence of its host in an animal infection model *in vivo*. These seemingly contradictory results have only been observed in one other mycovirus system^26^ to our knowledge. In most mycovirus infections, effects on *in vitro* growth are correlated with effects on host virulence, with both overall positive and overall negative impacts being described^18,19,27^. Although rare, increased fungal virulence due to mycovirus infection could lead to increased spread and transmission of both host and mycovirus^12,28^, depending on lifetime output of zoospores. However, *in vitro* growth may not correlate with growth in the wild. It is possible that the specific culture media used contributes to substrate-specific negative impact of BdDV-1 on its host, as has been described with mycoviruses BbPMV-1 and BbPMV-3^29^. On the other hand, the relatively low zoospore loads observed on deceased frogs infected with v+ isolates may indicate that BdDV-1 negatively impacts *Bd* growth both *in vitro* and *in vivo*. If this is the case, then BdDV-1 may be altering *Bd* phenotype in a way that increases virulence without increasing host growth. This could be through the increased production of virulence factors, which in *Bd* are hypothesized to include metalloproteases and Crinkler Necrosis (CRN) genes^17,30,31^. We observed that *in vitro*, a strain with the endogenized virus (CLFT067) differentially expressed 4 M36 metalloproteases relative to virus negative strain, CLFT071. Surprisingly the M36 metalloproteases were all downregulated in the v+ strain relative to the v-. It is conceivable that the expression of metalloproteases during infection triggers the frog immune system, with downregulation associated with immune evasion. Resolving this issue will require understanding the expression of the M36 metalloproteases and other virulence-associated genes, which are known to have upregulated expression on the frog host^32^.

The broad pattern of vertical inheritance of BdDV-1 in *Bd* lineages may relate to its integration in the fungal chromosome. Rather than a means of genome replication, the endogenization of CRESS viruses is thought to be “accidentally” mediated by the Rep protein during viral replication in the host nucleus^33^. If BdDV-1 inserted itself in the genome early in the origin of *Bd*, it has been maintained as a non-pseudogenized, expressed cluster of genes located in a dynamic region of the genome– subtelomeric and between a high copy number locus (ribosomal RNA) and a DNA integrase. Our PCR results targeting the flanks of the viral integration locus indicate that all viral positive *Bd*-BRAZIL strains share the same integration locus. The co-evolution of *Bd* and BdDV-1 and shared integration locus suggest a single integration event and subsequent vertical transmission of BdDV-1. Although BdDV-1 may be primarily expressed from a chromosomal location and inherited through the fungal DNA, its presence is variable across *Bd*-BRAZIL isolates, and its nuclear location would not limit its ability to cause a profound effect on host phenotype. For example, the hypovirulence caused by CHV1 of *C. parasitica* can be recapitulated by expression from an integrated cDNA version of the virus^34^. Moreover, CRESS and other viruses are capable of maintaining both sequence conservation and gene expression following endogenization^35,36^. Additional studies are underway to develop a transfection system to provide a more controlled environment to test the phenotypic effects of BdDV-1.

The demonstration that *Bd* harbors CRESS viruses opens up the potential for exploiting their unique biology for use in genetic engineering and biocontrol of the fungus. A DNA virus of chytrid fungi may allow the development of viral-based vectors to genetically transform chytrids, a much needed research tool. CRESS viruses also have potential to be used as a form of sprayed biocontrol as the ssDNA viruses may show environmental transmission^19,37^. Before any efforts to undertake viral-based remediation or biocontrol strategies, it is imperative to know more about the distribution and potential impact of BdDV-1 and related viruses across amphibian hosts and their parasites. Pointedly, in the case of BdDV-1, our results show an association of the virus with increased rather than decreased fungal virulence, making its application in this area unlikely. Instead, the *Bd*-mycovirus system may reinforce the emerging notion that hyperparasites can optimize host virulence to increase their Darwinian fitness^38^.

In conclusion, we describe a novel CRESS virus discovered in *Bd* that is primarily present as endogenous copies in the genomes of enzootic lineages and which may influence the growth and virulence of the host. Because of this phylogenetic pattern, we hypothesize that defense against or escape from a natural enemy such as BdDV-1 could be related to the emergence of *Bd*-GPL as the dominant genotype across the world. The deep phylogenetic history of the virus, which is found at an even higher prevalence in CAPE & ASIA1 clades sister to the GPL + BRAZIL lineages, suggests that the common ancestor of most *Bd* lineages would have been infected by BdDV-1 with losses of the virus common in the *Bd*-BRAZIL lineage and before the global dispersal of *Bd*-GPL. By comparing naturally infected *Bd*-BRAZIL isolates with uninfected isolates, we showed the virus is associated with increased virulence, though the experimental infection and *in vitro* conditions provide only a small snapshot of the true environmental conditions experienced by *Bd*. Finally, BdDV-1 occurs in the wild in regions in which active hybridization of *Bd* is occurring^4,39^. The threat that hybridization could bring BdDV-1 into a GPL background and increase the virulence of hybrid genotypes is a serious concern. In fact, experimental trials have shown that *Bd* hybrid strains (between *Bd*-GPL and *Bd*-BRAZIL) could be more virulent than parental lineages when infecting Brazilian and African frog hosts^40,41^. Viral spillover from hybrids into *Bd-*GPL could have severe implications for amphibian population declines by increasing the virulence of already hypervirulent *Bd* lineages.

## Acknowledgements

LKF-L, TYJ, JES are Fellows in CIFAR program Fungal Kingdom: Threats and Opportunities. The work was partially supported by a catalyst grant from CIFAR and CIFAR fellowship funds. The Gordon and Betty Moore Foundation Award #9337 (<u>10.37807/GBMF9337</u>) to LKF-L, TYJ, and JES supported RC, EK, and MY. LKF-L also was supported by a Pew Scholar award from the Pew Charitable Trusts. LFT was granted by the São Paulo Research Foundation (FAPESP #2016/25358-3) and the National Council for Scientific and Technological Development (CNPq #302834/2020-6).

## Author contributions

Conceptualization, R.C., M.Y., T.Y.J, J.E.S.; methodology, R.C., M.Y., T.C., T.Y.J, L.K.F, J.E.S.; formal analysis, R.C., M.Y., T.Y.J, L.K.F, J.E.S.; investigation, T.C., E.F., E.K.; access to samples and sequencing data, T.S.J, L.F.T., T.Y.J; writing – original draft, R.C., M.Y., T.Y.J, J.E.S.; writing – review & editing, L.K.F.-L., E.K., T.C., D.R.S., L.F.T.; data and microscopy visualization: M.Y., E.K., R.C., D.R.S., T.C., L.K.F.-L.; supervision, T.YJ., J.E.S., L.K.F.-L.; project administration, T.Y.J., J.E.S.; funding acquisition, L.K.F.-L., L.F.T., T.Y.J., J.E.S.

## Resource availability

### Lead contact

Further information and requests for resources and reagents should be directed to and will be fulfilled by the lead contact, Jason Stajich.

### Materials availability

No unique reagents were generated in this study.

### Data and code availability

- Genome sequence and annotation for CLFT044 and CLFT071 have been deposited at DDBJ/ENA/GenBank. The primary sequence data for Nanopore and Illumina DNA and RNA sequence have been deposited as NCBI BioProjects. *Bd-*BRAZIL strain sequence data have been deposited under an NCBI BioProject and individual strain data has been deposited to the NCBI Sequence Read Archive. All accession numbers are listed in the key resources table and all data are publicly available as of the date of publication.
- All original code for phylogenetic analyses have been deposited at github and is publicly available as of the date of publication. DOIs are listed in the key resources table.
- Any additional information required to reanalyze the data reported in this paper is available from the lead contact upon request.

## Experimental model and subject details

### Culture Acquisition

All *Bd* isolates are deposited in CZEUM (Collection of Zoosporic Eufungi at the University of Michigan)^65^ and maintained on 1% tryptone (1%T) growth medium at room temperature (approximately 21° C). All isolates used belong to the non-GPL *Bd*-BRAZIL lineage and were genetically characterized^66^. Five isolates were determined to be virus positive through genome sequence analysis (CLFT061, CLFT067, CLFT044, CLFT148, CLFT139), and four additional isolates were determined to be virus negative (CLFT068, CLFT070, CLFT071, CLFT144).

### Amphibian infection model

We performed all investigations involving live vertebrate animals following published protocols^41^ approved by the University of Michigan Institutional Animal Care and Use Committee (IACUC protocol PRO00009614). Adult Dwarf Clawed frogs (*Hymenochirus boettgeri*) were acquired from vendors Live Aquaria and VWR. Upon arrival, the frogs were group-housed in 37 liter aquaria at a maximum of 20 frogs per tank. After a 3-5 day acclimation period, the frogs were heat treated to remove any incoming *Bd* infection following previous work^72^. The tank temperature was raised 1 °C per day until reaching a maximum temperature of 30 °C. The tanks were maintained at 30 °C for 7 days, and then lowered by 1 °C per day until returning to 21 °C ambient room temperature. Frogs were not sexed prior to the infection experiment and were randomly sorted into treatment groups.

## Method details

### Viral discovery from screening *Bd* sequences

Published isolate DNA sequence Illumina reads were downloaded from the National Center for Biotechnology Information (NCBI) Sequence Read Archive (SRA) and additional *Bd*-BRAZIL strains (**Table S1, S3**). Illumina sequence reads were aligned to the *Bd* JEL423 genome with bwa v0.7.17^42^ and processed with samtools to identify those reads which did not align to the genome (samtools fastq -f 4 BAMFILE). BdDV-1 genomic coverage from the Illumina sequence reads was calculated using mosdepth v0.3.3^43^. BdDV-1 coverage and *Bd* phylogeny visualizations were made using ggtree v3.8.2^44^. These reads were further assembled with SPAdes v3.15.2^45^. The resulting contigs were screened by translated searches against UniProt databases which identified regions of assembled contigs as homologous to viral *Rep* genes. The ORFs were further predicted from the contig assembly with prodigal v2.6.3^46^.

### CRESS Rep phylogeny

Virus Rep proteins from representatives of the major families of CRESS viruses (*Geminiviridae*, *Genomoviridae*, *Circoviridae*) were obtained from the UniProt database. The Rep proteins from these viruses and BdDV-1 were aligned with MUSCLE v5.1^47^, and a phylogeny was constructed with IQTREE v2.2.1^48^ using 1000 bootstrap replicates. Phylogenetic tree visualization was rendered with ggtree v3.8.2^44^. Software scripts for these analyses are archived at https://github.com/myacoub005/BdDV1_Discovery; DOI: 10.5281/zenodo.10662592. These methods were repeated to include the two Rep-like proteins from BdDV-1 in the phylogeny to assess their relationship to the primary Rep protein.

### Strain phylogeny

A phylogeny of the *Bd* strains was constructed from raw alignments of the reads to JEL423 genome following GATK best practices and assuming diploid genomes ^49,50^. A NJ phylogenetic tree of the strains constructed with Poppr v2.9.3^51^. Software and scripts to run the variant calling, mapping of reads, detection of unmapped reads, and tree construction are archived in https://github.com/stajichlab/PopGenomics_Bd; DOI: 10.5281/zenodo.10662979.

### BdDV-1 tanglegram

The *Cap* gene nucleotide sequences from the Illumina genomes were aligned using MUSCLE v5.1 and the phylogeny was constructed with IQTREE v2.2.1^48^ with 1000 bootstrap replicates. A tanglegram was constructed against the *Bd* strain phylogeny using the R package dendextend v1.17.1^52^.

### Illumina assembly of strain CLFT044

The genomes of strain CLFT044 and additional *Bd*-BRAZIL strains **(Table S1)** were sequenced using paired-end (2×125 bp) sequencing on an Illumina NextSeq HiSeq-2500 v4 platform. Reads were trimmed for adapters and quality using Trimmomatic^53^ and *de novo* assembled with SPAdes v3.15.5^45^. After identification of a single contig of 3,647 bp in CLFT044 that contained blast matches to the *Rep* gene, the putative viral contig was compared to itself using the BLAST algorithm implemented in YASS v1.16^54^. We also inspected the assembly graph produced by spades for evidence of loops indicating circularity using the software Bandage v0.8.1^55^.

### RNA extraction and sequencing of CLFT044

RNA was extracted from 7 day old cultures of CLFT044, CLFT067 and CLFT071 grown on 1% Tryptone Agar at 18 °C using TRIzol (Invitrogen, Mulgrave, VIC, Australia) following manufacturer’s instructions and incorporating a 12 hour isopropanol precipitation to increase yields. Total RNA was sent to *Novogene* (Davis, CA) for polyA enrichment and Illumina NovaSeq PE 2×150 sequencing to obtain 6G raw data per sample. Reads from CLFT044 were aligned to the assembled CLFT044 genome using bwa v0.7.17^42^ to determine transcriptional activity of BdDV-1.

Reads from CLFT067 (v+) and CLFT071 (v-) were aligned against the JEL423 reference genome concatenated with the BdDV-1 gene using the R package kallisto v.0.48.0^56^. Differential expression analysis was performed using the R package DESEQ2 v.1.40.2^57^ with a minimum log2fold change > 2 and Bonferroni adjusted p-value < 0.05.

### Nanopore assembly and annotation of *Bd*-BRAZIL

Tissue was collected from 7 day old cultures of CLFT044 and CLFT071 grown on 1% tryptone agar at 21 °C by flooding plates with 1 mL of sterile RO water, scraping colonies with an L-spreader, and collecting into a 2 mL sterile tube. Tubes were centrifuged at 6500 rcf to remove supernatant and flash frozen with liquid nitrogen. DNA was extracted from tissue using a Cetyltrimethylammonium Bromide (CTAB) extraction protocol^58^. The DNA was sent to MiGS (now SeqCenter; Pittsburgh, PA, United States) to obtain 900 Mb (∼30X coverage) of Oxford Nanopore sequence reads for CLFT044 and CLFT071. The reads were *de novo* assembled using Canu v2.2^59^ followed by 10 iterations of polishing with Pilon v1.24^60^ with Illumina sequence data. Annotation was performed using Funnanotate v1.8.14^61^ which trained *de novo* gene predictors using RNAseq data combine with alignment of homologous proteins to achieve high annotation accuracy.

### Identification of the viral locus

The BdDV-1 locus was identified through BLASTP v2.13.0+^62^ searches using the viral *Rep* gene obtained from the Illumina *Bd* genomes against the CLFT044 ONT genome annotation. The raw Illumina FASTQ data from v+ and v- strains were aligned against the CLFT044 ONT genome using bwa v0.7.17^42^ to identify the full length 4.405 kb region that comprises BdDV-1. This region was covered only by v+ strains and absent in all v- strains. The 4 proteins in this region were searched against the UniProt and NCBI databases to confirm they were viral in origin.

### Genomic integration comparison

Scaffold_10, possessing BdDV-1 in *Bd*-BRAZIL strain CLFT044, was compared to the homologous scaffolds in virus-negative *Bd*-BRAZIL strain CLFT071 and *Bd*-GPL strain JEL423. Homologous scaffolds were identified using minimap2 v2.24^63^ with scaffold_10 as the query against both virus-negative assemblies. We generated the scaffold comparison visualizations with gggenomes v0.9.9.9000^64^ using the results from minimap2 to plot linkages between assemblies.

### Tests of viral presence and circularity

Viral presence was confirmed using PCR with BdDV-1 specific primers to amplify a 591 bp fragment. The circularity of the virus was confirmed on a subset of samples by using 2 BdDV-1 specific primer sets that were reverse complements of one another. One primer set amplified a ∼800 bp region, and the other set amplified the remaining ∼1400 bp of the 2.2kb circular viral genome. See **Table S2** for primer sequences. All PCR reactions followed the manufacturer recommended recipe for Phusion High Fidelity Polymerase (New England BioLabs, Inc.) with these PCR conditions: 98 °C 30 s, 40 cycles (98 °C 10 s, 60 °C 20 s, 72 °C 30 s), 72 °C 7 min.

Rolling circle amplification (RCA) was attempted on a subset of samples that were confirmed to be virus positive through PCR testing. A TempliPhi 100 Amplification Kit (Cytiva) was used, following a slightly modified protocol^23^. BdV-2F and BdV-3R PCR primers were added to the reaction to target BdDV-1. RCA products were digested with a *Sca*I restriction enzyme (Thermo Fisher Scientific) following manufacturer instructions. Additional attempts were made using only a single virus specific primer, following manufacturer protocol, and using the restriction enzyme *Dra*I.

### Fluorescence in situ hybridization (FISH)

#### Probe synthesis and labeling

Polynucleotides were synthesized by PCR using Phusion High-Fidelity DNA polymerase (NEB) and locus-specific primers (see **Table S2** for sequences). To target the BdDV-1 genome, a 2143 bp region was amplified from CLFT044 total DNA. For the *Bd* actin locus, a 2058 bp probe was synthesized from JEL423 total DNA.

Following spin column purification (Macherey-Nagel), the PCR products were directly labeled with Alexa Fluor 555 dye and purified using the FISH Tag DNA Orange Kit (ThermoFisher Scientific). Labeling was performed according to the manufacturer’s instructions, with the following modification: 0.95 µL DNase I working solution and 1.2 µg of template DNA were used in the nick translation reaction. Labeled probe concentrations were determined spectrophotometrically using a NanoDrop 2000 (Thermo Scientific).

#### Cell fixation and staining

*Bd* strains JEL423 and CLFT044 were grown in 1% tryptone (w/v) at 24 °C in tissue culture-treated flasks (Fisher), washed once and resuspended in Bonner’s salts (10.27 mM NaCl, 10.06 mM KCl, 2.7 mM CaCl_2_ in MilliQ water) to a final concentration of 1×10^7^ zoospores/mL. 96-well glass bottom plates (Eppendorf) were plasma cleaned, immediately coated with a 1 mg/mL Concanavalin A (Sigma) solution for 8 min, washed thrice with Bonner’s salts, and overlaid with 120 µL of the resuspended *Bd* zoospores. Cells were allowed to adhere to the Concanavalin A for 5 min before fixation with 200 µL of 4% paraformaldehyde in 50 mM cacodylate buffer (pH 7.2). Cells were fixed on ice for 20 min, then for 10 min at room temperature before being washed three times with PBS (137 mM NaCl, 2.7 mM KCl, 10 mM Na_2_HPO_4_, 1.8 mM KH_2_PO_4_) for 5 min. Samples were then permeabilized in 0.2 % Triton X-100 in PBS for 12 min at room temperature (about 23 °C) and washed thrice with PBS.

Cells were then rinsed once with prehybridization buffer (50 % deionized formamide [Invitrogen], 2xSSC [0.3 M NaCl, 0.03 M sodium citrate, pH 7.0], 10 % dextran sulfate [Sigma-Aldrich]). 75 µL prehybridization buffer was then added to each well, the plate was sealed with adhesive aluminum foil and incubated for 30 min at 37 °C and subsequently transferred to a preheated lab oven set at 85 °C for 12.5 min to denature cellular DNA. During this time, FISH probes were prepared by diluting labeled probes in prehybridization buffer to a final concentration of 1.5 ng/µL, denatured at 93 °C for 5 min, placed on ice for 1 min, and then kept at room temperature (about 25 °C).

Following oven incubation, the 96 well plate was briefly centrifuged, the prehybridization buffer was gently aspirated with a micropipette and 75 µL denatured probe mix was added to each well. Immediately afterwards, the plate was resealed, moved to a humid hybridization chamber, and incubated for 38-42 h at 37 °C. Unbound probes were removed by low-stringency washes at room temperature. Specifically, cells were rinsed briefly once with 2xSSC, twice for 5 min in 2xSSC, twice in 0.4xSSC for 5 min, once for 5 min in 4xSSC, twice in PBS and then counterstained with 1µg/mL DAPI in PBS for 10 min, rinsed twice with PBS, and then overlaid with PBS for imaging.

#### Microscopy and image analysis

Cells were imaged on a Nikon Ti2-E inverted microscope equipped with a 100x oil PlanApo objective and sCMOS 4mp camera (PCO Panda). Images were acquired at room temperature using both DIC microscopy, and epifluorescence microscopy with excitation light at 400 nm to visualize DAPI and 550 nm to visualize the Alexa Fluor 555 dye-conjugated probes. Images were further processed in Fiji^67^. The number of hybridization signal spots per cell was determined from overlays of DAPI and FISH probe micrographs by manual counting. A total of 4-9 pictures were analyzed for each sample. Final micrographs for Figure 1E were composed in Inkscape (Inkscape Project, 2020).

### Confirmation of viral integration

Viral integration was confirmed using PCR primers to amplify the 1023 bp region spanning the right flank of the integration locus: *Rep* gene (BdVlf-F) to *Bd* ITS1_SSU region (BdVlf-R). See **Table S2** for primer sequences.

### Curing attempts

Virus positive isolates were streaked onto 1% tryptone plates. After 3-4 days of growth single zoosporangia were isolated onto fresh 1%T plates with a sterile wire tool. The virus positive isolates were also grown on 0.1 mg/L cycloheximide 1%T plates. After streaking, cultures onto media with cycloheximide, single zoosporangia were transferred to fresh 0.1 mg/L cycloheximide 1%T plates. After 3-4 days of growth, single zoosporangia were isolated to fresh 1%T plates. The same method was used with 80 µM ribavirin instead of cycloheximide. All single zoosporangia isolations were allowed to grow to saturation then tested for viral presence by PCR.

### Transmission electron microscopy

Isolates CLFT044 v+ and CLFT068 v- were grown to saturation on 1%T plates. Zoospores were harvested from the plates by flooding them with 1 mL of sterile water. Zoospores were chemically fixed, infiltrated, and embedded for transmission electron microscopy^68^. Ultrathin serial sections of zoospores were examined and imaged on a JEOL JEM-1400 at the University of Michigan Microscopy Core, Ann Arbor, MI, USA.

### *In vitro* growth assays

Four virus positive and four virus negative isolates were grown independently on 1%T plates. After 7 days, each plate was flooded with 2 mL of liquid 1%T media and left to incubate at room temperature (approximately 21° C) for 30 min. Zoospores were then harvested from each plate. The zoospore concentration of each isolate was estimated using a Multisizer 4e Coulter Counter (Beckman Coulter, Inc.).

For the first assay, each isolate was diluted into 1 mL of liquid 1%T media at a concentration of 10^6^ zoospores/mL in a 24-well plate with 2 replicates per isolate. The isolates were allowed to grow at room temperature (approximately 21° C) for 5 days, and the optical density (OD) was recorded every hour using a Epoch 2 Microplate Spectrophotometer (BioTek Instruments).

For the second assay, zoospore dilutions were prepared as described above, except in a total volume of 0.8 mL 1%T media in a 48-well plate with 5 replicates per isolate. Only 6 of the isolates were used (3 virus positive and 3 virus negative). The lid of the plate was treated with Triton X-100^69^ to prevent condensation on the underside of the lid. The isolates were allowed to grow for 12 days with an OD measurement every hour. The plates were agitated by gentle shaking for 5 seconds before each OD measurement. The third assay was prepared as described for the second assay.

### qPCR for *Bd* and BdDV-1

Real-time quantitative PCR was performed on DNA extracted swabs to determine *Bd* zoospore equivalents (ZSE)^71^. Swabs from control frogs and frogs infected with virus negative isolates of *Bd* were single-plexed using only a *Bd* specific qPCR assay. Swabs from frogs infected with virus positive isolates of *Bd* were multiplexed using both a *Bd* specific assay and a BdDV-1 specific assay. See **Table S2** for primer and probe sequences. Standards for viral copy number were created by cloning PCR product using custom BdDV-1 primers into the pCR 2.1-TOPO vector using the TOPO TA cloning kit (Invitrogen). Transformed *Escherichia coli* colonies with successful insert were plasmid extracted using the Zyppy Plasmid Miniprep kit (Zymo Research). Plasmid copy numbers were estimated using the formula:

Ten-fold dilutions of viral standards were created with 10^6^ plasmid copies through 10^0^ plasmid copies. Ten-fold dilutions of *Bd* standards were created starting with DNA extracted from 10^6^ zoospore suspensions. All qPCR reactions were run on a QuantStudio 3 qPCR machine (Applied Biosystems).

A growth assay was performed using qPCR to quantify *Bd* zoospore equivalents and BdDV-1 viral copies through the growth period. The growth assay was set up as described above, using 12-well plates with 2 mL working volumes and 4 replicates per isolate. Initial concentrations of each isolate were normalized to a 0.001 OD reading instead of a zoospore count. Starting after 1 day and measuring every 3 days for a 16 day growth period, 100 μL of each replicate was removed and added to a microcentrifuge tube containing 400 μL of 100% ethanol and stored at −20 °C. To prepare for extraction, each sample was spun down at 13,000 rcf for 10 minutes, then all liquid was removed by pipette. Samples were then DNA extracted using Prepman^TM^ Ultra (Applied Biosystems) following manufacturer protocols. qPCR for *Bd* zoospores and BdDV-1 viral copies was performed as described above.

### Experimental infection

To prepare inoculum, zoospores were harvested and concentrations estimated for each of the 8 *Bd* isolates using a Coulter counter as described above for the growth assays. However, instead of flooding with liquid 1%T media, plates were flooded instead with sterile water. The zoospore suspensions were diluted to a concentration of 2.5×10^5^ zoospores/mL in autoclave sterilized RO water with aquarium salt. 10 mL of each suspension were aliquoted into sterile 50 ml conical tubes.

Post-heat treatment, 90 frogs were chosen at random and sorted into randomized experimental groups with 10 frogs per group. Each frog was weighed as a proxy for body size, and then immediately placed in an individual 50 mL tube with the zoospore suspension. Ten frogs were placed into 50 mL tubes with *Bd*-free autoclave sterilized RO water with aquarium salt as a control group. The frogs remained in the tubes for a 6 hour inoculation, then the frog and the entire zoospore suspension were added to individual 0.9 L aquaria.

The frogs were housed and cared for in the individual aquaria for the remainder of the 60 day experiment with daily monitoring for mortality and morbidity. Frogs were removed from the tanks and swabbed using Medical Wire & Equipment fine tip rayon swabs at day 10, day 30, day 60, and at death or euthanasia. Swabs were DNA extracted using a DNeasy blood and tissue kit (Qiagen). Living frogs were weighed at day 30 and day 60 to estimate health and body condition.

## Quantification and statistical analysis

A Baker’s Gamma Correlation Coefficient was calculated to determine the congruency of the *Bd* genome and the BdDV-1 *Cap* phylogenies using the R v4.3.1 package dendextend v1.17.1. The correlation coefficient was calculated as 0.95, suggesting congruent phylogenies between fungus and virus.

RNAseq transcripts from three replicates of CLFT067 (v+) and CLFT071 (v-) were aligned against the JEL423 reference genome concatenated with the BdDV-1 genes using the R package kallisto v0.48.0^56^. Differential expression analysis was performed using the R v4.3.1 package DESEQ2 v.1.40.2^57^ with a minimum log2fold change ≥ 2 and Bonferroni adjusted p-value < 0.05.

The results of all statistical analyses described here are present in the Results section. For all spectrophotometer *in vitro* growth assays, efficiency of growth was measured as the difference between the average OD at the final three time points and the average OD at the first three time points. The growth rate was determined using the means of the highest positive slopes during the growth period^70^. For each assay, a nested t-test was performed in GraphPad Prism v10.1.0 (https://www.graphpad.com/) to determine differences in the efficiency of growth or growth rate between virus positive and virus negative isolates.

To account for the differences in ITS copy numbers between *Bd* strains, the zoospore equivalents determined by qPCR assay for each strain were adjusted relative to the strain with the highest ITS copy number. We assessed the ITS copy number of each *Bd* strain by aligning the raw Illumina FASTQ data for each strain against the 18S and 28S sequences adjacent to BdDV-1 in the CLFT044 assembly using bwa v0.7.17^42^. We then divided the 18S and 28S coverage by the overall genome coverage to estimate the ITS copy number. After this estimation, we adjusted the qPCR results of each strain by dividing by the ratio of the stain ITS copy number to the ITS copy number of the highest strain (CLFT148 v+) to correct for ITS variation. A nested t-test was performed in GraphPad Prism to determine if there was a difference between average zoospore load on dead frogs infected with virus positive or virus negative *Bd* isolates.

A Cox proportional hazards model, clustered by isolate, was used to determine differences in hazard ratios between v+ and v-infected frogs. A Spearman correlation test was performed to determine if initial body weight influenced the probability of survival of *Bd*-infected frogs. These tests were performed in R v 4.3.1.

