## Supplementary figures and images for "Discovery of an endogenous DNA virus in the amphibian killing fungus and its association with pathogen genotype and virulence"

### Supplemental Figure 1

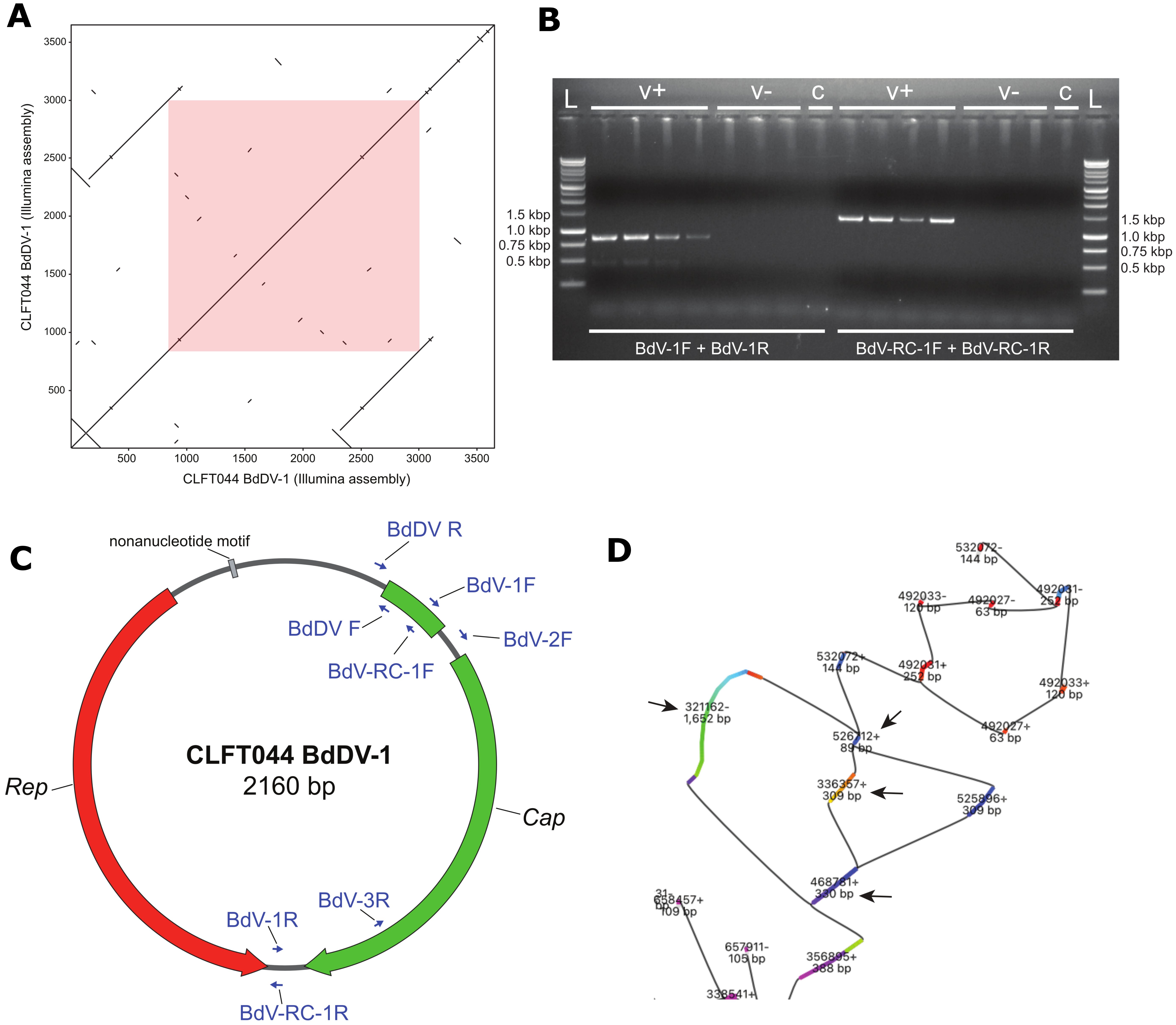

### Supplemental Figure 2

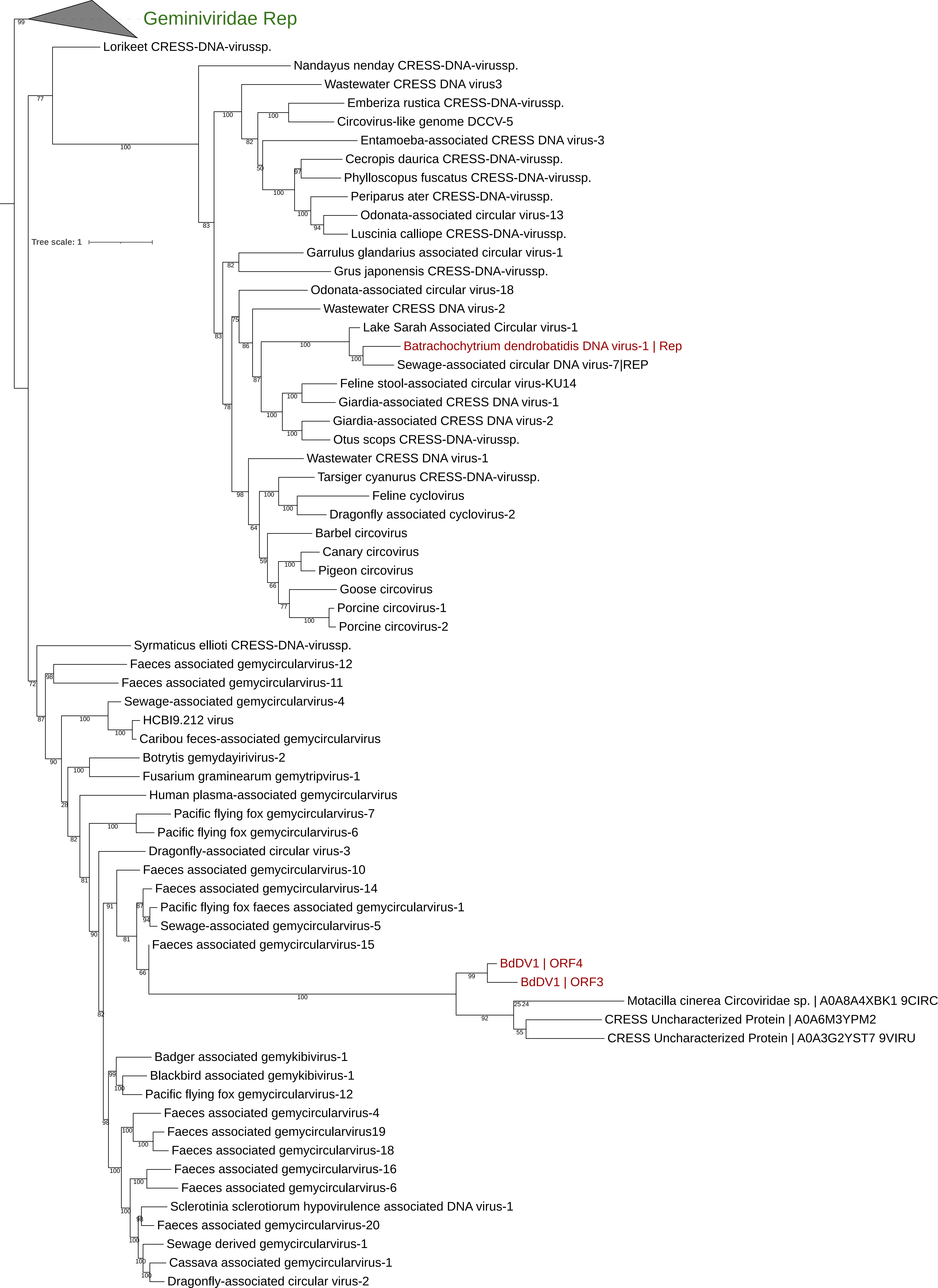

### Supplemental Figure 3

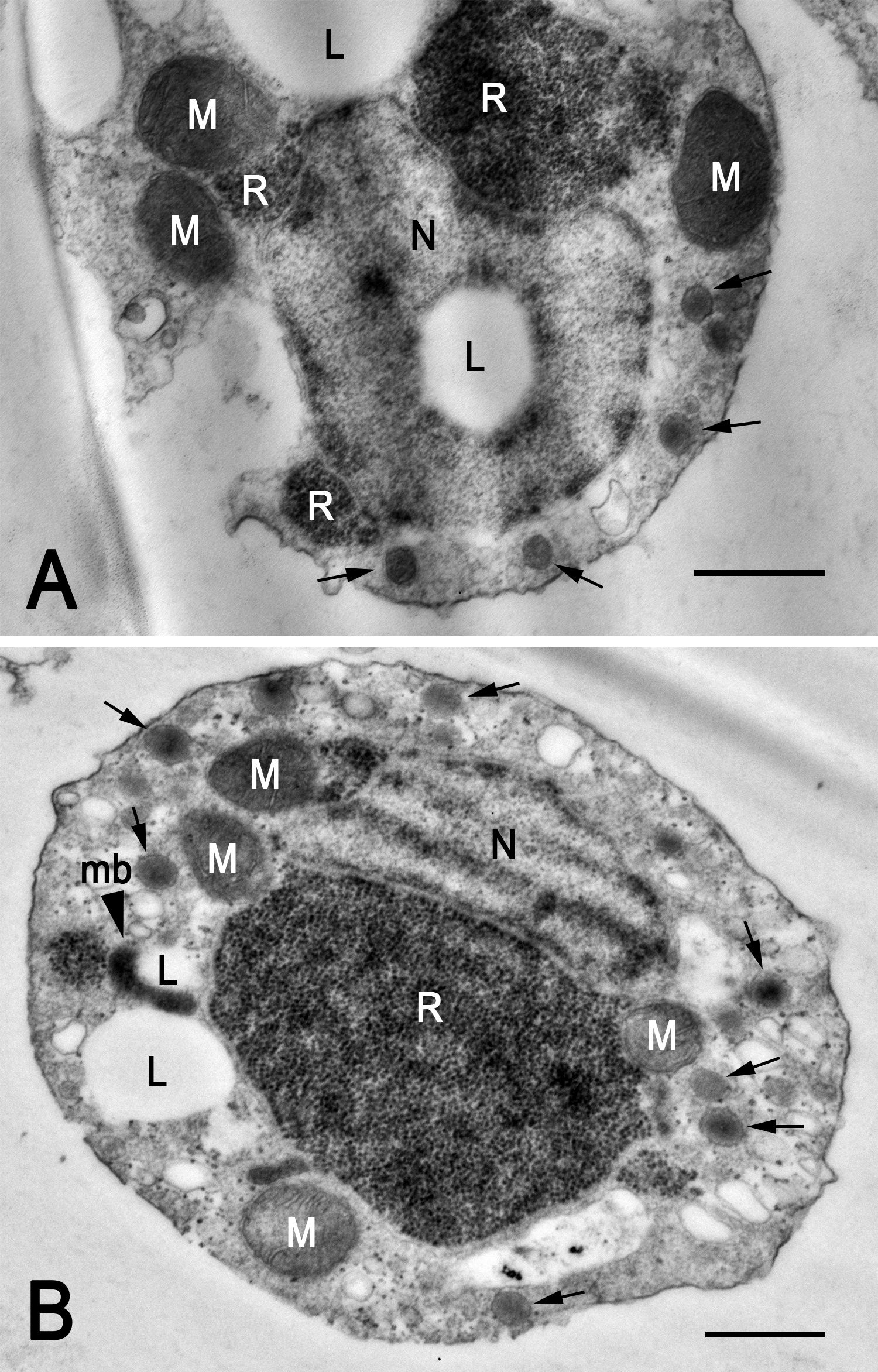

### Supplemental Figure 4

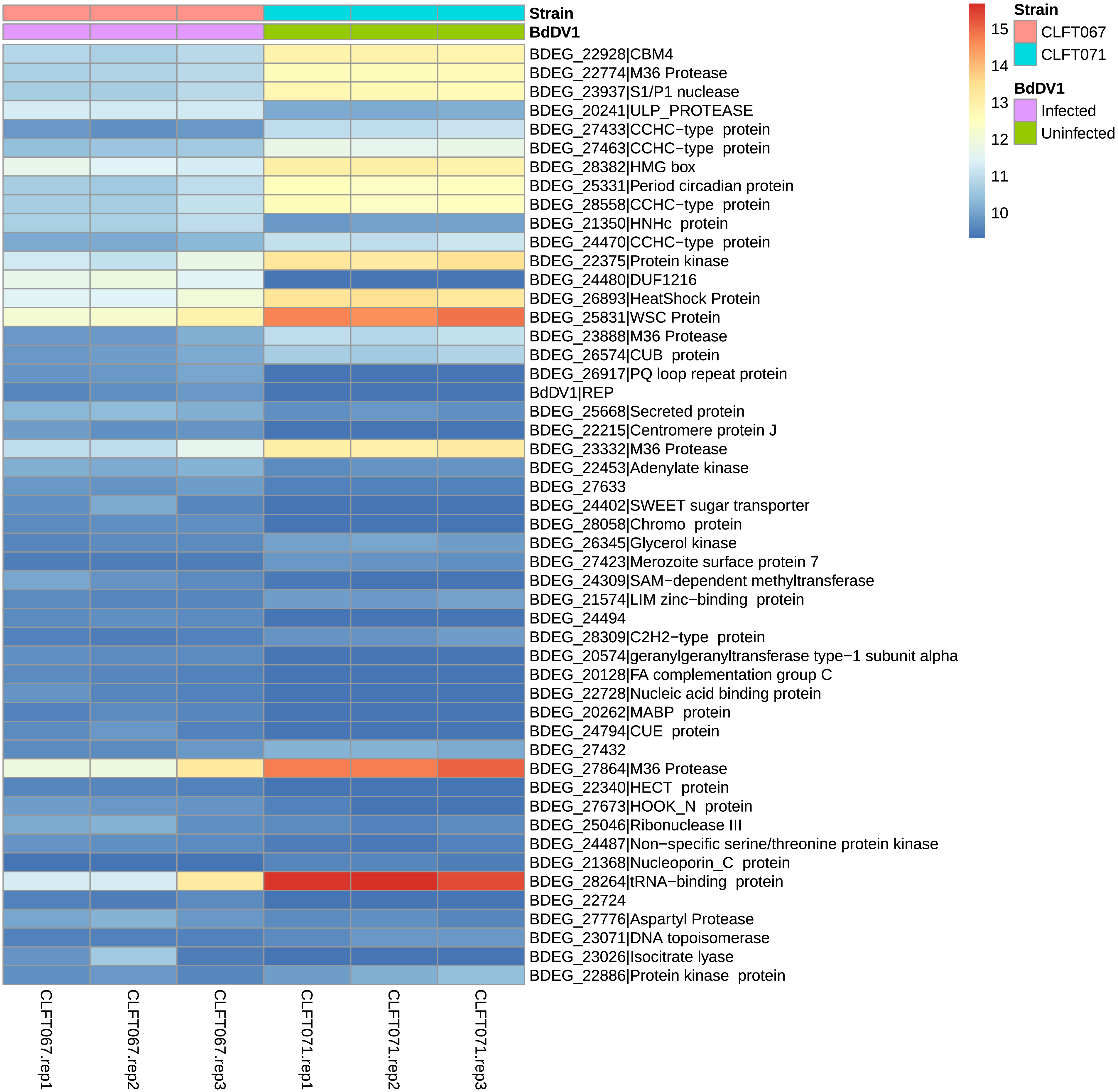

### Supplemental Figure 5

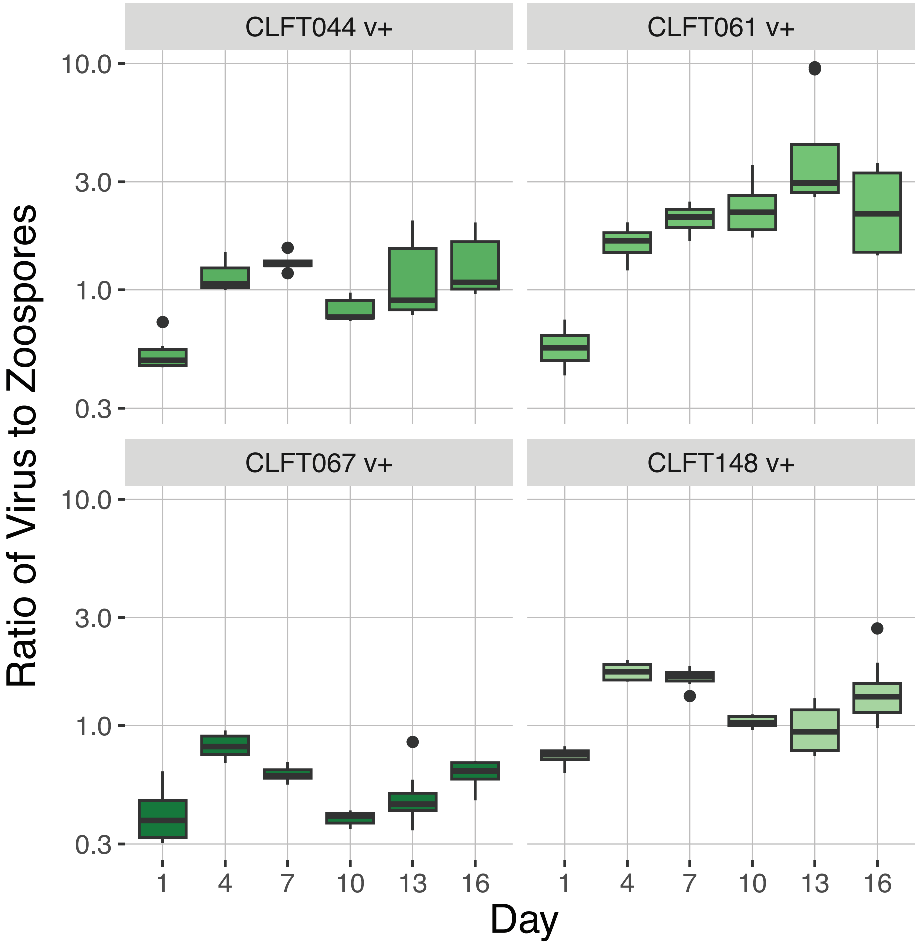

### Supplemental Figure 6

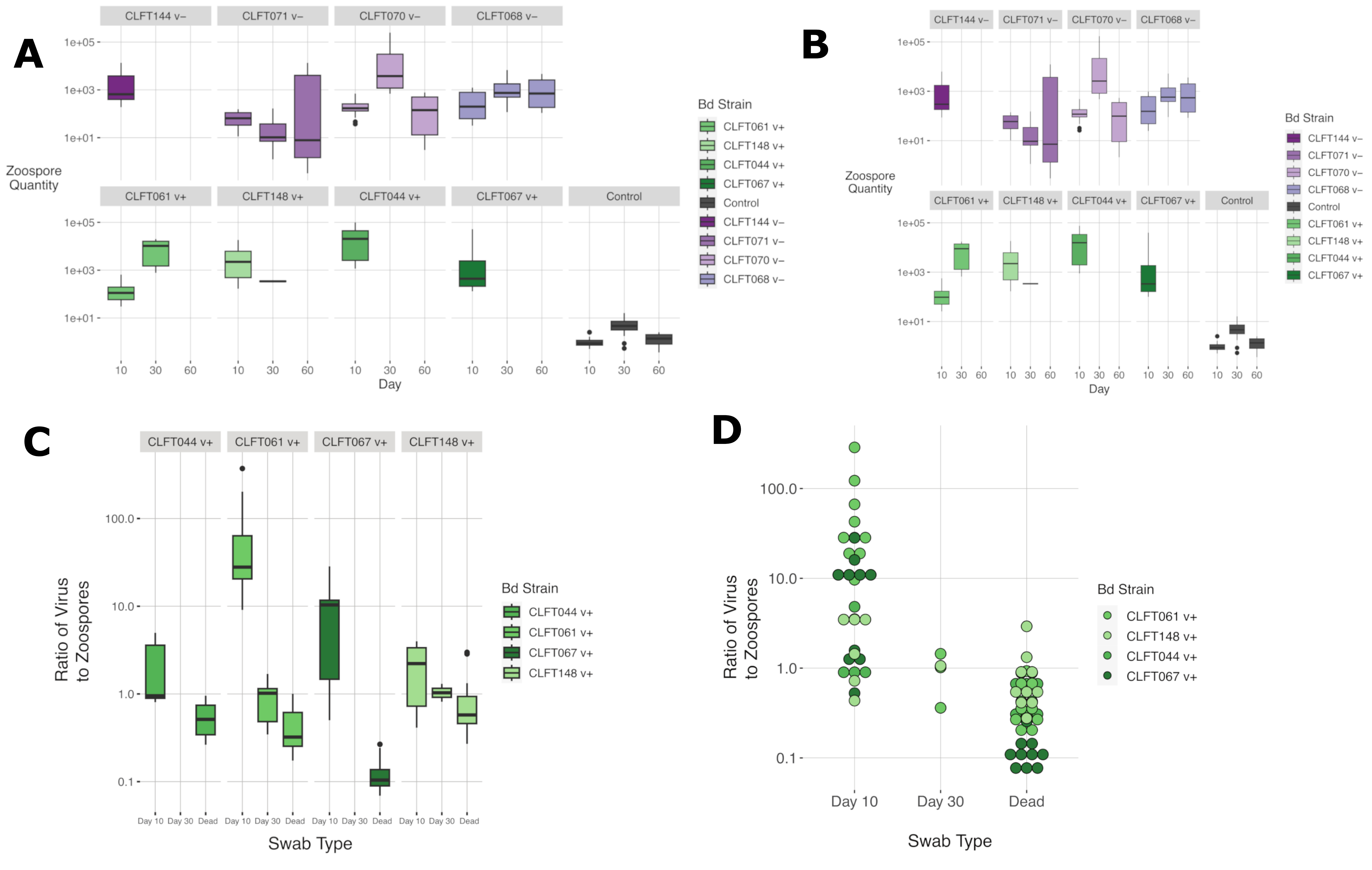
