## Supplemental Tables 1 and 2 for "Discovery of an endogenous DNA virus in the amphibian killing fungus and its association with pathogen genotype and virulence"

**Supplemental Table 1.** *Bd* isolates used in this study presenting collection information and viral presence. Brazilian state abbreviations are Paraná (PR), Santa Catarina (SC), and São Paulo (SP). Isolates indicated with asterisks are those whose genomes were resequenced for this study.

| **Isolate** | **Lineage** | **Year** | **Geographic origin/ Municipality** | **State** | **Host Species** | **Collector** | **Viral Presence** |
|---|---|---|---|---|---|---|---|
| CLFT044* | *Bd*-BRAZIL | 2013 | Serra da Graciosa, Morretes | PR | *Hylodes cardosoi* | C.M. Betancourt | Positive |
| CLFT061 | *Bd*-BRAZIL | 2013 | Pomerode | SC | *Hylodes meridionalis* | C.M. Betancourt | Positive |
| CLFT067 | *Bd*-BRAZIL | 2013 | Serra do Japi, Jundiaí | SP | *Hylodes japi* | C.M. Betancourt | Positive |
| CLFT139* | *Bd*-BRAZIL | 2014 | Serra da Graciosa, Morretes | PR | *Hylodes cardosoi* | T.S. Jenkinson | Positive |
| CLFT148* | *Bd*-BRAZIL | 2014 | Serra da Graciosa, Morretes | PR | *Hylodes cardosoi* | T.S. Jenkinson | Positive |
| CLFT068* | *Bd*-BRAZIL | 2013 | Serra do Japi, Jundiaí | SP | *Hylodes japi* | C.M Betancourt | Negative |
| CLFT070* | *Bd*-BRAZIL | 2013 | Serra do Japi, Jundiaí | SP | *Hylodes japi* | J.E. Longcore | Negative |
| CLFT071 | *Bd*-BRAZIL | 2013 | Serra do Japi, Jundiaí | SP | *Hylodes japi* | C.M. Betancourt | Negative |
| CLFT144 | *Bd*-BRAZIL | 2014 | Serra da Graciosa, Morretes | PR | *Hylodes cardosoi* | T.S. Jenkinson | Negative |

**Supplemental Table 2**: Primer and probe sequences used for viral presence confirmation, viral circularity confirmation, DNA FISH, viral integration confirmation, and viral qPCR.

| | |
|---|---|
| **Virus Confirmation** | |
| BdV-2F | GTCAGAATTTGACGGGGGTA |
| BdV-3R | CCGACAACAATTTGCAACAG |
| **Viral Circularity** | |
| BdV-1F | CACTGCTTTTCCGTCAACAA |
| BdV-1R | CAAGGGTCCTACTGGACCAA |
| BdV-RC-1F | TTGTTGACGGAAAAGCAGTG |
| BdV-RC-1R | TTGGTCCAGTAGGACCCTTG |
| **Viral Integration** | |
| BdVIf-F | AAGCGCTATCACTGGCTTCG |
| BdVIf-R | AGAATTCGTGTGCACCCTCC |
| **DNA FISH probes** | |
| BdActin-F | TGTTCACGCCCACTGCTATT |
| BdActin-R | AGGGCCAGACTCGTCATACT |
| BdDV-2F | GGTGATTGGTAAAAATTTGAGCTGT |
| BdDV-2R | TACAAGACAAGGAGTTTGGTGG |
| **Virus qPCR** | |
| BdDV F | CCTGAGTACCCTGATCACAATGT |
| BdDV R | GGGTCATTGGTCGTATCTTCA |
| BdDV Probe | MBGNFQ-CCATGGTGGCGTTCT-NED |
